# An invariant network architecture with broadly expressed regulators distinguishes cell-type from cell-state identity *in situ*

**DOI:** 10.64898/2026.10.25.761612

**Authors:** Rui Zhou, Li Ge, Jing Li, Zeqian Xu, Min Chen, Yan Guo, Daniel M. Czajkowsky, Zhifeng Shao

**Affiliations:** State Key Laboratory of Systems Medicine for Cancer, School of Biomedical Engineering, Shanghai Jiao Tong University, Shanghai 200240, China; College of Computing and Data Science, Nanyang Technological University, Singapore 639798, Singapore

## Abstract

Core regulatory networks (CRNs) of tissue-specific master transcription factors (TFs) are essential for defining cell identity and are widely used to reprogram cells and to decipher pathogenic mechanisms. However, to date, CRNs have only been described at the cell-type level: their characteristics in the distinct states that a single cell type can adopt have not been examined. This leaves open what the cell-type CRNs actually represent. Here, using the zonated hepatocyte states of the mouse liver as a model system, we construct the first state-level CRNs in a native tissue. For this purpose, we developed a robust LCM-Spatial-ATAC-seq method that achieves bulk-level data quality from microdissected cell populations. By identifying what is shared among the state-level CRNs, we determined what is cell-type defining: an invariant sub-network of core-regulatory TFs (crTFs) that maintains identical positions and conserved interactions in the CRNs. Unexpectedly, the top positions of this invariant sub-network are occupied by broadly expressed TFs rather than the well-established master TFs, which reside downstream. Yet the state-level CRNs are clearly different, both in terms of composition and interactions. The former are dominated by zone-specific regulators, whereas the latter are spread throughout the network, encompassing about one third of all interactions between the states. Thus, our results reveal a novel dual character of a cell’s CRN: an invariant sub-network that defines the cell-type identity, together with a variable peripheral network that underlies its state specialization.

## Introduction

Each of the hundreds of cell types in a multicellular organism maintains a distinct functional identity. Foremost among the mechanisms that establish this identity is the core regulatory network (CRN), composed of cross-regulating transcription factors (crTFs) that effectively acts as the "engine" driving the expression program underlying a cell’s characteristics (Neph et al. 2012; Wilkinson et al. 2017; Jung and del Sol 2020; Almeida et al. 2021). These networks have been defined through the TFs with the most cell-type-specific expression or super-enhancer association, often considered the "master regulators" of cell identity (Zhou et al. 2017; D’Alessio et al. 2015), or as highly interconnected sub-networks of super-enhancer-associated TFs (Saint-André et al. 2016; Hnisz et al. 2013; Whyte et al. 2013). As such, CRNs are central to the fundamental question of what distinguishes one cell type from another (Neph et al. 2012) and are increasingly applied in mechanistic studies of disease development (Jung and del Sol 2020; Zhang et al. 2023) and in efforts to reprogram cell types for replacement strategies (Ieda et al. 2010; Vierbuchen et al. 2010; Du et al. 2014).

Yet CRNs have to date been defined only at the level of (at most) cell types: how they change among the distinct, context-dependent cell states that cells often adopt within any given cell type (Trapnell 2015; Rafelski and Theriot 2024) remains unknown. Indeed, many metazoan cell types exist in multiple well-defined, stable states in tissues in response to their local micro-environment, such as the intestinal enterocytes along the crypt-villus axis and dermal fibroblasts in the papillary and reticular layers of the skin (Moor et al. 2018; Beumer et al. 2022; Philippeos et al. 2018). Hence, with this in mind, for CRNs determined at the cell-type level, it is actually not clear what such networks represent. In practice, these networks are typically constructed from samples such as bulk tissues, in which the identity and homogeneity of the constituent cells are assumed rather than established (Jung and del Sol 2020; Almeida et al. 2021; Saint-André et al. 2016). A type-level CRN is thus, in effect, a population-wide average over these states, with no clear understanding of how it relates to the CRN of any one state. At one extreme, cell states may reflect the transcriptomic response to extrinsic signals, with the CRN itself essentially remaining intact; the type-level network would then represent every state. At the other, the CRN may be significantly reorganized between states, making the type-level network a composite of state-specific components. Does such reorganization involve additional master regulators? Is it modular or integrated throughout the network? Answers to these questions are unknown but would provide insight into both how state specialization is established and what is preserved to maintain cell-type identity.

The features shared among the established states of a cell type may thus provide a more reliable basis for defining its cell-type CRN. The liver offers an excellent system to delineate this idea. Within each liver lobule, hepatocytes exist in different cell states as a result of their proximity to the local vasculature, a phenomenon known as liver zonation (Gebhardt 1992; Jungermann and Keitzmann 1996; Ben-Moshe and Itzkovitz 2019). Single-cell RNA-seq analysis has revealed that more than half of the expressed genes in hepatocytes differ substantially between the cell states (Halpern et al. 2017). There are also well-established molecular markers for the different hepatocyte states (Halpern et al. 2017), and the layered cellular architecture of the liver lobules is advantageous for investigations of cells within tissue sections (Cunningham and Porat-Shliom 2021). To date, the hepatocyte CRN has been defined from bulk liver tissues, comprising a core circuitry of 13 TFs and an extended network of 26 additional TFs (Dubois-Chevalier et al. 2023). Although restricted to the best-characterized factors and thus unlikely to represent the complete CRN, this network provides a useful reference for state-level comparisons.

Here, we developed a robust LCM-Spatial-ATAC-seq method, enabling ATAC-seq acquisition with laser capture microdissection (LCM) under immunofluorescence guidance, to profile chromatin accessibility in the three zonated hepatocyte states of the mouse liver lobule. Combining these profiles with super-enhancer annotation, chromatin interaction (Hi-C) maps, transcriptomic data, and TF binding motif analysis (Saint-André et al. 2016), we constructed the first state-level CRNs in a native tissue context. We found that the three CRNs share an invariant sub-network of 143 crTFs, with identical positions and conserved interactions. This sub-network defines hepatocyte cell-type identity. Unexpectedly, its top positions in the sub-network hierarchy are occupied by broadly expressed factors, with the established master regulators (D’Alessio et al. 2015; Whyte et al. 2013) residing at mid-level hub positions. Beyond this invariant sub-network, the CRN is indeed reorganized between states. There are changes in composition, involving a few tens of crTFs that are dominated by zone-specific regulators rather than additional master regulators, together with integrated rewiring of ∼36% of interactions between the states. Finally, we find that the core circuitry of the previously defined hepatocyte CRN largely overlaps with the invariant sub-network, whereas over half of its extended-network factors are not invariant crTFs (Dubois-Chevalier et al. 2023), a result only apparent from state-level characterization. Overall, these results show that a cell’s CRN comprises two distinct components: an invariant sub-network of crTFs that maintains cell-type identity, and a variable peripheral network whose reorganization establishes cell-state specialization, with implications for reprogramming and disease.

## Results

### LCM-Spatial-ATAC-seq achieves bulk-level data quality from spatially defined cell populations

For an effective spatial ATAC-seq method using LCM on tissue sections, two critical technical requirements must be met: the enzymatic reactions must be highly efficient for cells within the tissue environment, and the native cellular and genomic architecture must be maintained throughout the process. Existing spatial ATAC-seq approaches perform the Tn5 tagmentation reaction on tissue sections attached to solid slides (Carraro et al. 2023; Deng et al. 2022; Llorens-Bobadilla et al. 2023; Russell et al. 2024), which makes optimization particularly challenging: different sections contain variable cell numbers, resulting in inconsistent Tn5-to-cell/DNA ratios, and the slide creates a physical barrier that also limits enzyme access to the nuclei proximal to the glass (Fig. 1a). We reasoned that all ATAC-seq steps should instead be performed on free-floating sections in the same tube before immunostaining and microdissection (Liu et al. 2026) so that precise calibration of the Tn5-to-cell/DNA ratio across all sections could be achieved with efficient reagent access to the nuclei from all sides (Fig. 1a).

**Fig. 1.**
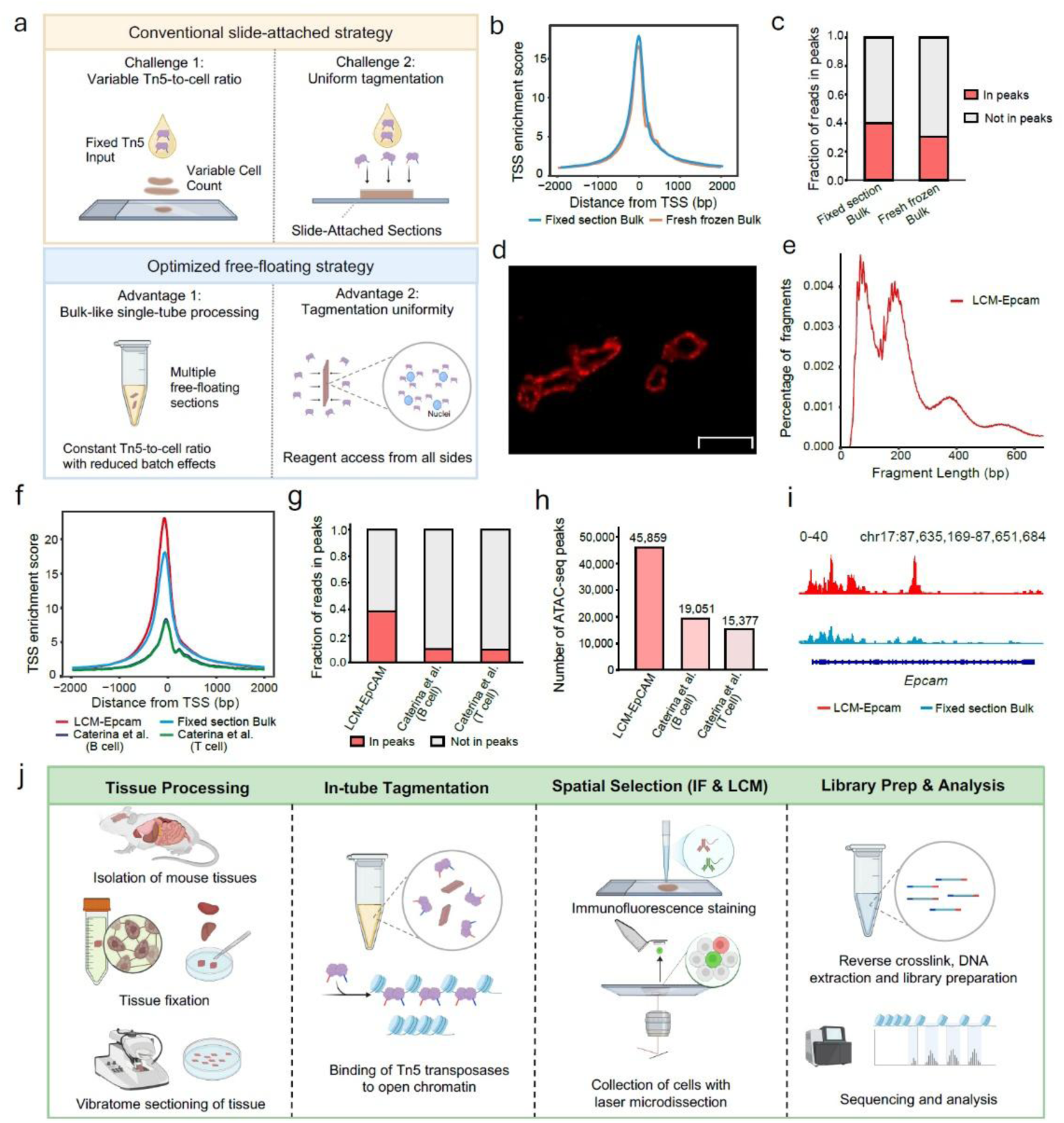
LCM-Spatial-ATAC design, workflow and data quality. **a** Schematic illustration comparing the technical considerations of the conventional slide-attached strategy (top) and the optimized free-floating strategy (bottom) of spatial ATAC-seq for tissue sections. Created with BioRender.com. **b** Comparison of the enrichment of reads at TSSs between merged fixed-section bulk ATAC-seq data from mouse kidney tissue and published merged fresh-frozen bulk ATAC-seq data from mouse kidney tissue (Liu et al. 2019). TSS, transcription start site. **c** Comparison of the FRiP scores between the ATAC-seq data generated from merged fixed-section bulk ATAC-seq data of kidney tissue and published merged fresh-frozen bulk ATAC-seq data of kidney tissue. FRiP, fraction of reads in peaks. **d** A typical immunofluorescence image of the epithelial cells in a kidney tissue section, labelled with an anti-EpCAM antibody. Scale bar: 50 μm. **e** Fragment length distribution of LCM-captured EpCAM+ (LCM-EpCAM) ATAC-seq data. **f** Comparison of TSS enrichment profiles among LCM-EpCAM ATAC-seq data, published LCM-captured B cell and T cell ATAC-seq data (Carraro et al. 2023), and merged fixed-section bulk ATAC-seq data from mouse kidney tissue. LCM, laser capture microdissection; bp, base pairs. **g** Comparison of FRiP scores among LCM-EpCAM ATAC-seq data and published LCM-captured B cell and T cell ATAC-seq data (Carraro et al. 2023). **h** Number of peaks identified for each sample in (g). **i** Genome-browser tracks showing ATAC-seq signals at the Epcam locus in LCM-EpCAM (red) and fixed section bulk kidney ATAC-seq (cyan) datasets. For a fair comparison of signal intensity, bigWig tracks were normalized to the same number of valid reads and displayed on the same y-axis scale. **j** Schematic workflow for spatially resolved chromatin accessibility profiling using LCM-Spatial-ATAC-seq. Created with BioRender.com.

Tissue sections also require chemical fixation to preserve structural integrity during the lengthy procedure, but over-fixation can hinder nuclear genome accessibility. After optimizing fixation conditions using cultured cells and then mouse kidney tissue sections (Methods, Supplemental Tables S1-S2), we found that 1% PFA fixation combined with our modified protocol yielded data surpassing the ENCODE ATAC-seq standards, with transcription start site (TSS) enrichment scores of ∼14 and fraction of reads in peaks (FRiP) scores of ∼40% (ENCODE Project Consortium, https://www.encodeproject.org/atac-seq/). These values are comparable to published results from bulk ATAC-seq with unfixed frozen kidney tissues (Fig. 1b, c, Supplemental Fig. S1), and substantially exceed those reported for prior LCM-ATAC-seq methods (Carraro et al. 2023) (TSS enrichment of 1 to 9, FRiP of 10 to 22%).

We then incorporated immunofluorescence labeling and LCM. As an initial test, we labeled epithelial cells in mouse kidney sections with anti-EpCAM antibodies (Fig. 1d) and performed LCM dissection of the labeled populations. The resulting data exhibited the expected nucleosomal and subnucleosomal patterns (Fig. 1e) and again surpassed ENCODE standards (Fig. 1f-h, Supplemental Fig. S1), with an estimated average yield of ∼5,000 unique fragments per cell. Fragments at the epithelial marker EpCAM locus were substantially enriched compared with bulk kidney ATAC-seq profiles, as expected from the selection of EpCAM-positive cells (Fig. 1i). Altogether, these results demonstrate that LCM-Spatial-ATAC-seq can indeed provide bulk-level data quality from localized cell populations. Given this per-cell yield, we collected ∼4,000 cells per replicate to ensure the depth required for robust CRN construction.

### State-resolved chromatin accessibility within the intact liver lobule

To construct state-level CRNs within a native tissue, we focused on the mouse liver lobule, where hepatocytes exist in three spatially defined, well-characterized metabolic states along the periportal-to-pericentral axis (Gebhardt 1992; Jungermann and Keitzmann 1996; Ben-Moshe and Itzkovitz 2019). Using LCM-Spatial-ATAC-seq (Fig. 1j), we profiled chromatin accessibility in these three major hepatocyte states: periportal (PV) hepatocytes adjacent to the portal vein, mid-zonal (MZ) hepatocytes, and pericentral (CV) hepatocytes surrounding the central vein (Fig. 2a). We labeled PV cells with an anti-E-cadherin antibody and CV cells with an anti-glutamine synthetase antibody, and identified MZ cells by their defined spatial position between the PV and CV-labeled regions (Fig. 2a, Methods).

**Fig. 2.**
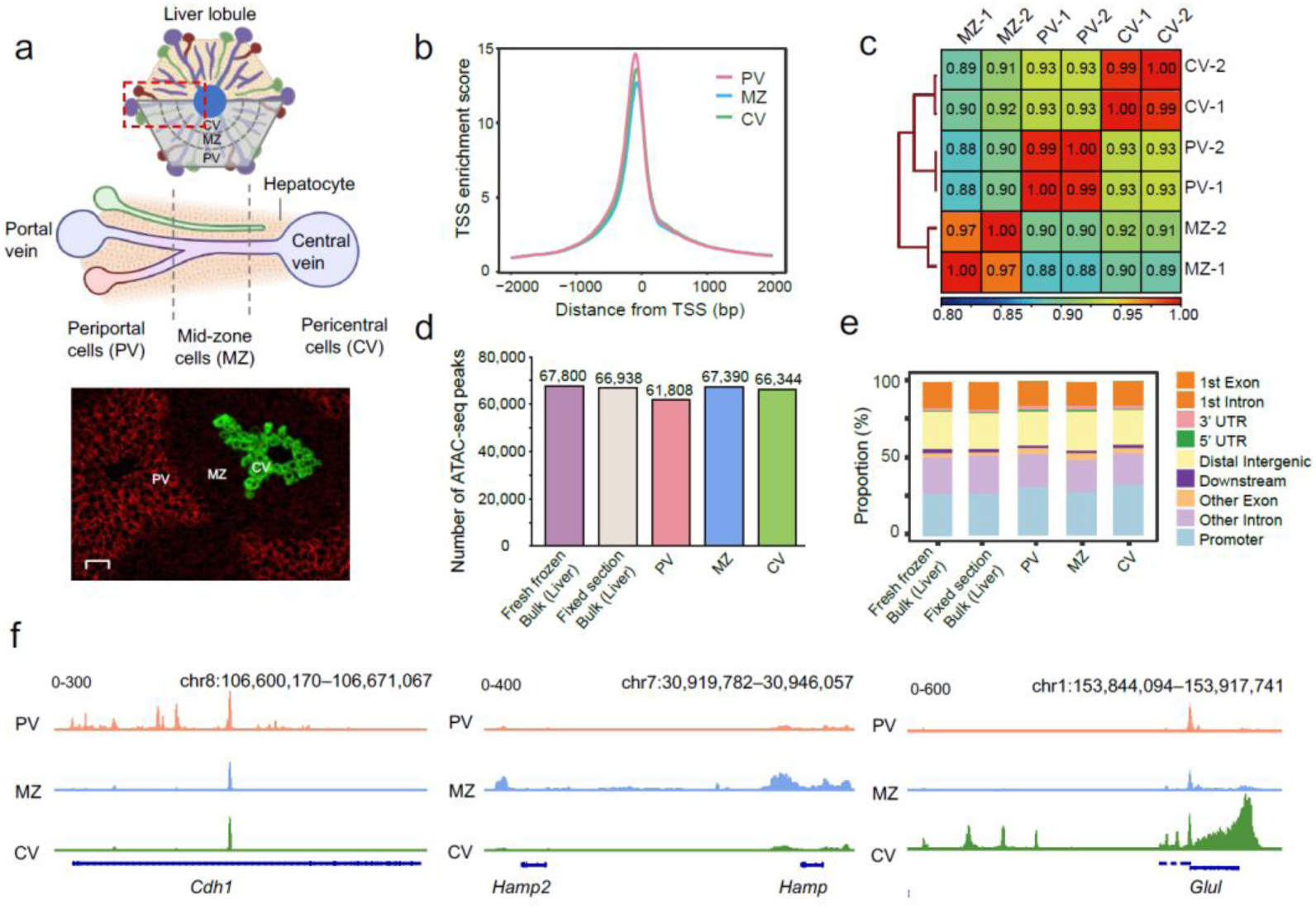
Spatially resolved chromatin accessibility in zonated hepatocytes revealed by LCM-Spatial-ATAC-seq. **a** Schematic representation of liver lobule organization showing the spatial arrangement of hepatocytes along the periportal-to-pericentral axis, including periportal (PV), mid-zonal (MZ), and pericentral (CV) hepatocytes, together with a representative immunofluorescence image used for anatomical identification of hepatocyte states. PV hepatocytes were labeled with an anti-E-cadherin antibody (red), CV hepatocytes were labeled with an anti-glutamine synthetase antibody (green) and MZ hepatocytes were defined by their spatial position between the PV- and CV-labeled regions. Scale bar, 50 μm. Created with BioRender.com. **b** Enrichment of reads at TSSs in merged ATAC-seq data from biological replicates of PV, MZ, and CV hepatocytes. **c** Heatmap of all-by-all Pearson correlations among PV, MZ and CV hepatocyte ATAC-seq datasets. Correlations were calculated using read counts across genome-wide 1,000 bp bins. **d** Comparison of the number of peaks identified in data derived from LCM-captured PV, MZ, and CV hepatocytes with fresh frozen and fixed section bulk liver ATAC-seq datasets. For PV, MZ, and CV groups, peaks were identified from the merged data of two independent biological replicates. Fresh frozen bulk liver ATAC-seq reference data were obtained from (Liu et al. 2019). **e** Genomic annotation of ATAC-seq peaks for each sample in (d). **f** Genome-browser tracks showing ATAC-seq signals at representative state-associated marker loci in PV, MZ, and CV hepatocytes. BigWig tracks were normalized to the same number of valid reads.

We obtained two independent biological replicates (∼4,000 cells per replicate) for each state. The replicate datasets exhibited high quality (Fig. 2b and Supplemental Fig. S2a, b) and high reproducibility, clustering by spatial zone (Fig. 2c). After pooling valid fragments from replicates for each state, peak calling identified ∼62,000-67,000 peaks per state (Fig. 2d, e and Supplemental Table S3), consistent with previous bulk ATAC-seq profiling of liver nuclei (Liu et al. 2019). State-specific marker genes exhibited both promoter and enhancer peaks (Fig. 2f). The ability to clearly resolve distal enhancer elements across all three states was essential for the subsequent reconstruction of state-specific CRNs. To our knowledge, these data represent the first in situ, spatially resolved chromatin accessibility landscapes of defined cellular states within an intact mammalian tissue.

To assess the extent to which chromatin accessibility differs between hepatocyte states, we compared the PV and CV peak landscapes, focusing on these two states because their marker genes and zonation functions are the best characterized (Ben-Moshe and Itzkovitz 2019; Halpern et al. 2017). Of the consensus peaks (present in both replicates) in each state, 40,045 were shared between PV and CV, while 7,585 were specific to PV and 12,490 to CV. Notably, 13,572 of 57,691 merged peaks (23.5%) were differentially accessible between PV and CV (|log2 fold change| > 1, consistent direction in both replicates), with 9,551 enriched in CV and 4,021 enriched in PV. The substantial fraction of differentially accessible peaks indicates that state specialization is accompanied by widespread remodeling of the chromatin landscape, extending well beyond a handful of marker loci.

To validate that these state-specific accessibility differences indeed reflect biological zonation, we examined established marker genes. CV marker genes (including Cyp2e1, Glul, Oat, and Cyp1a2) showed 3-to 39-fold higher accessibility in CV than PV, while PV marker genes (including Cyp2f2, Sds, Hal, and Ass1) showed 3- to 14-fold higher accessibility in PV (Supplemental Fig. S3a). Importantly, Cdh1 (E-cadherin), the antigen used for PV immunolabeling, and Glul (glutamine synthetase), the antigen used for CV immunolabeling, both showed the expected zone-specific accessibility, validating the dissection strategy. Differentially accessible peaks were significantly enriched near marker genes of the corresponding state (odds ratio = 18.6 for CV, 10.2 for PV; Fisher’s exact test, p < 10^-30^), and PV-enriched peaks were absent near CV marker genes. Furthermore, changes in chromatin accessibility correlated strongly with changes in gene expression among differentially expressed genes (Pearson R = 0.66, p < 10^-16^; Supplemental Fig. S3b). While this fold-change agreement demonstrates that the differential accessibility reflects transcriptional differences, we note that promoter accessibility alone is not deterministic of expression levels, which depend significantly on distal enhancer activity (Klemm et al. 2019). These results confirm that the state-resolved accessibility landscapes faithfully capture biological zonation and provide a reliable basis for the construction of state-specific CRNs.

### State-resolved CRNs reveal a conserved crTF composition across hepatocyte states

A CRN is defined as the interaction network of robustly expressed TFs that cross-regulate with each other, termed crTFs (Jung and del Sol 2020; Su et al. 2022). A defining feature of crTFs is that they are regulated by super-enhancers (SEs), which are thought to ensure their stable expression. We define CRNs in our data according to two criteria (Fig. 3): crTFs are expressed and associated with SEs active in the vast majority of cells in a given state, and crTFs exhibit cross-regulatory interactions with each other.

**Fig. 3.**
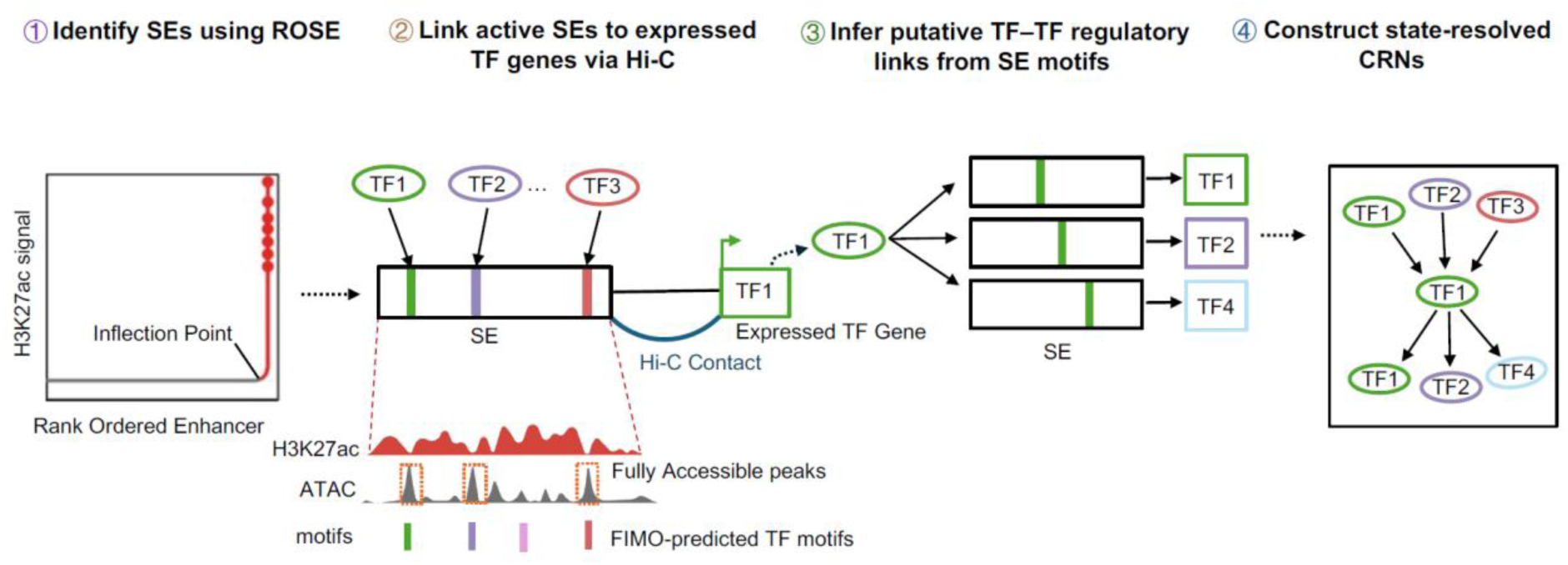
Computational pipeline for identifying core regulatory networks (CRNs). Schematic representation of the computational pipeline for CRN identification. Super-enhancers (SEs) were identified from H3K27ac ChIP-seq data using ROSE. State-active SEs were defined as SEs overlapping at least one highly accessible ATAC-seq peak, which was identified using a randomized distribution with a probability score > 0.9. State-active SEs were linked to expressed transcription factor (TF) genes (CPM > 1) using significant long-range chromatin interactions from Hi-C maps. FIMO was then used to identify predicted TF-binding motifs within highly accessible peaks in state-active SEs. Motif-containing SEs linked to expressed TF genes were used to infer putative TF–TF regulatory interactions and construct state-resolved CRNs. ROSE, Rank Ordering of Super-Enhancers; SE, super-enhancer; TF, transcription factor; Hi-C, high-throughput chromosome conformation capture.

To satisfy the first criterion, we identified SEs in hepatocytes by applying ROSE (Lovén et al. 2013) to H3K27ac ChIP-seq data (Supplemental Fig. S4a and Supplemental Table S3). State-active SEs were defined as those overlapping at least one highly accessible ATAC-seq peak, identified using a probability score > 0.9 based on a randomized distribution (Methods, Supplemental Fig. S4b). Using this approach, ∼98% of all SEs were found to harbor at least one highly accessible peak in each individual state (Supplemental Fig. S4c and Supplemental Table S3). We then identified TF genes expressed in each state (CPM > 1, Methods, Supplemental Table S4) and linked state-active SEs to these genes using statistically significant long-range chromatin contacts from our previously published mouse liver Hi-C maps (Liu et al. 2026), rather than linear proximity alone (Saint-André et al. 2016; Huang et al. 2018). These SE-TF associations showed a many-to-many relationship across hepatocyte states (Supplemental Fig. S4d, e and Supplemental Table S4), yielding 478 unique SE-associated TFs in PV, 520 in MZ, and 545 in CV hepatocytes (Supplemental Table S4).

For the second criterion, we used FIMO (Grant et al. 2011) to identify which SE-associated TFs had known binding motifs within highly accessible peaks in SEs linked to other SE-associated TF genes, defining putative cross-regulatory interactions (Fig. 3). Overall, we identified approximately 200 crTFs in each state (Supplemental Table S5). Notably, this set contains not only tissue-specific regulators but also a significant number of broadly expressed TFs that satisfy all criteria for core regulatory function. Their inclusion follows directly from our criteria, which only require super-enhancer association and cross-regulation but impose no tissue-specificity filter. Thus, we find an intricately wired regulatory landscape, comprising thousands of interconnected interactions in each state, defining the state-resolved CRNs for PV, MZ, and CV hepatocytes (Supplemental Table S5).

As a first step towards identifying the properties of these CRNs that are cell-type defining, we sought to determine the crTFs that are present in each CRN. We found that a significant majority, 180 of the 219 crTFs (82.2%), was shared among all three states, with the remaining crTFs roughly equally distributed between being present in only two states (n = 20) or only one state (n = 19) (Fig. 4a; Supplemental Table S6). As expected, several master regulators of hepatocytes are within the shared crTF set, including HNF4A, HNF1A, CEBPA, NR5A2 and FOXA2 (Dubois-Chevalier et al. 2023; Bravo González-Blas et al. 2024). Similarly, crTFs previously reported to function in only certain states, such as TCF7L1 and TBX3 (Bravo González-Blas et al. 2024), were found among the non-shared factors. Thus, the vast majority of crTFs that compose each state-specific CRN belong to a shared set that defines the hepatocyte cell type, although importantly, each CRN also includes a significant number of state-specific crTFs as well (Fig. 4a).

**Fig. 4.**
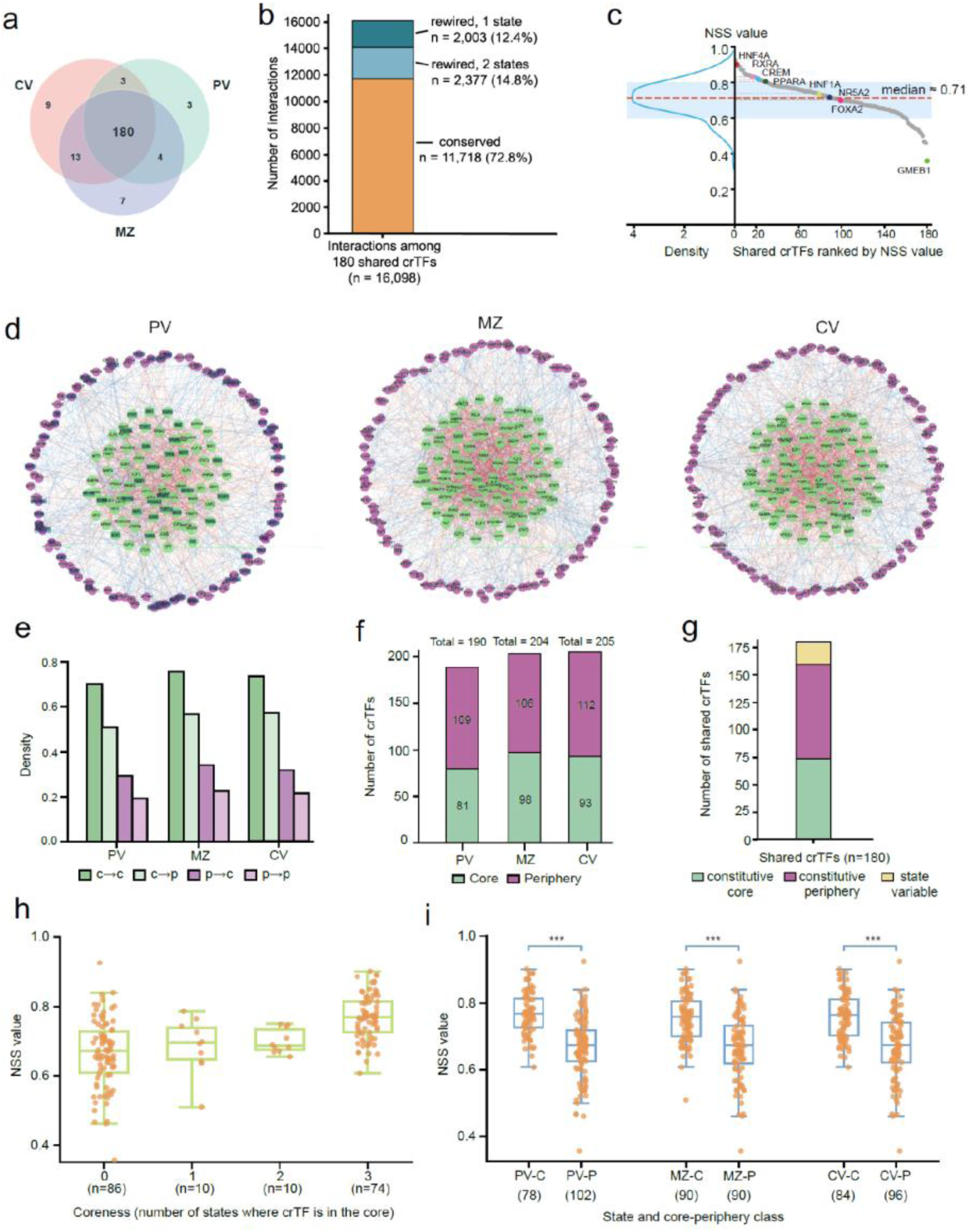
Network analysis of the state-resolved CRNs reveals both shared and state-specific features. **a** A Venn diagram showing the overlap of crTFs among PV, MZ, and CV hepatocyte CRNs. **b** Conservation of core regulatory interactions among the 180 shared crTFs. Of 16,098 interactions, 11,718 (72.8%) are conserved across all three states, 2,377 (14.8%) are present in two states, and 2,003 (12.4%) in a single state. **c** Distribution of Neighborhood Similarity Scores (NSS) for shared crTFs. NSS measures the proportion of conserved regulatory neighborhoods (Jaccard Index) across PV, MZ, and CV states (median 0.71). Selected key hepatocyte regulators are labeled. **d** Visualization of the core-periphery structure for the PV, MZ, and CV CRNs, partitioned using the Lip discrete core-periphery algorithm (p = 0.005 against randomized null networks); core crTFs (green) at the center, peripheral crTFs (purple) at the boundary. For visual clarity, 10% of edges were randomly sampled. **e** Edge density quantification within and between the core (c) and periphery (P) partitions. **f** Distribution of the total number of crTFs assigned to the core (green) and periphery (purple) partitions for each hepatocyte state. **g** Conservation of core-periphery assignments among the 180 shared crTFs across all states. Stacked bar represents the number of crTFs that are constitutive core (green, n = 74), constitutive periphery (purple, n = 86), or state variable (yellow, n = 20). **h** Spearman correlation between NSS values and coreness (number of states where a crTF is in the core) for shared crTFs (n = 180; ρ = 0.522, p = 7.03 x 10^-14^). **i** NSS distributions of shared crTFs assigned to the core (-C) versus periphery (-P) across zones, with group sizes (n) indicated below the x-axis. Significance levels are indicated above brackets (***p < 0.001, two-sided P values).

### Conserved interactions further define the shared CRN

Having established that the crTF composition is largely conserved across the states, we next examined the extent to which the interactions and organizational features of these networks are also conserved. To this end, we modeled each state-resolved CRN as a directed graph and analyzed the networks in terms of three basic levels of organization: pairwise interaction patterns, network topology, and regulatory hierarchy (Li et al. 2022; Rombach et al. 2017; Jothi et al. 2009).

At the first level, we found that ∼73% of interactions (11,718 of 16,098) among the 180 shared crTFs are conserved across all three states (Fig. 4b). These conserved interactions thus represent a second invariant feature of the hepatocyte cell-type CRN, beyond the shared crTF composition. The remaining ∼27% (4,380) are present in one state but absent in at least one other, indicating that even among the conserved regulators, a significant fraction of interactions are reconfigured between zones.

To quantify this rewiring for each shared crTF across the states, we calculated a neighborhood similarity score (NSS), defined as the Jaccard index of its state-specific neighborhoods in the PV, MZ, and CV states (Methods; Supplemental Table S6). Overall, NSS values ranged from 0.35 to 0.92 (median 0.71), with most (72.2%) exhibiting values between 0.60 and 0.80 (Fig. 4c). Thus, the neighborhoods of most of the shared TFs change by 20% to 40% between the states. Among the shared crTFs with the highest NSS (0.90) is HNF4A, often considered the master regulator of hepatocyte function (Kotulkar et al. 2023), indicating that its local regulatory neighborhood is exceptionally well-conserved between the states. Other key hepatocyte regulators also exhibited relatively high NSS values (Fig. 4c). At the opposite extreme, the shared crTF with the lowest NSS (0.35) is GMEB1, known to orchestrate glucose and lipid metabolic homeostasis in zonated hepatocytes (Thériault et al. 1999; Quagliarini et al. 2023), whose significant rewiring likely reflects the distinct metabolic programs of different hepatocyte states.

Interestingly, this analysis shows that no crTF exhibits a completely conserved set of interactions across the states: even the shared crTFs with the most conserved interaction neighborhoods undergo some degree of interaction rewiring between states. This partial conservation at the interaction level prompted us to ask whether higher-order organizational features remain invariant across the states.

### Conserved regulatory hierarchy and core-periphery structure define an invariant network architecture

We first tested whether each state CRN exhibits a modular community structure or a core-periphery structure (Rombach et al. 2017; Treviño et al. 2012). Community detection methods (Directed Louvain modularity, Infomap, OSLOM, and HSBM; Methods) yielded no statistically significant or biologically interpretable communities. In contrast, the Lip’s Discrete CP Partitioning algorithm (Lip 2011) revealed a significant core-periphery organization for each state (p = 0.005 against randomized null networks; Fig. 4d, e), with similar distributions of core and peripheral crTFs across PV, MZ, and CV (Fig. 4f and Supplemental Table S7).

Among the 180 shared crTFs, 160 (∼89%) maintained their core-periphery assignment across all states, including 74 constitutive core crTFs and 86 constitutive peripheral crTFs (Fig. 4g and Supplemental Table S7). Furthermore, shared crTFs with higher coreness tended to exhibit higher NSS values (Spearman ρ = 0.52; Fig. 4h), with core crTFs consistently showing higher NSS (∼0.76) than peripheral crTFs (∼0.67) in each state (Fig. 4i). Thus, core crTFs exhibited more conserved neighborhoods, while peripheral crTFs exhibited greater interaction variability. Non-shared crTFs were also frequently located in the periphery, further underscoring the higher variability of interactions in this region.

While the core-periphery analysis reveals the overall architecture, it does not distinguish the specific regulatory roles of individual crTFs (Jothi et al. 2009; Gerstein et al. 2012). We therefore characterized each crTF as occupying an upstream ("top"), mediator ("middle"), or downstream ("bottom") position based on its extent of outgoing and incoming interactions with other crTFs (Gerstein et al. 2012; Yu and Gerstein 2006).

The distribution of crTFs across these hierarchical levels was remarkably similar in all states: ∼16% to 19% in the top, ∼56% to 62% in the middle, and ∼22% to 25% in the bottom (Fig. 5a, b and Supplemental Table S8). This organization differed significantly from random networks (Supplemental Fig. S5), confirming that these CRNs are indeed hierarchical. Moreover, ∼89% of the shared crTFs maintained an identical hierarchical level across all three zones (Fig. 5c and Supplemental Table S8), indicating that the upstream-to-downstream regulatory roles are a robust property of the hepatocyte cell type. Combining this hierarchical conservation with the core-periphery conservation observed above, we identified 143 shared crTFs that maintain both an invariant structural positioning and a constant hierarchical level across all three states (Supplemental Table S9). We found that ∼74% (7,498 of 10,140) of the interactions of just these 143 shared crTFs are conserved across all states. Thus, this subset constitutes the invariant sub-network that defines the hepatocyte cell-type, providing a stable structural arrangement within which state-specific remodeling occurs.

**Fig. 5.**
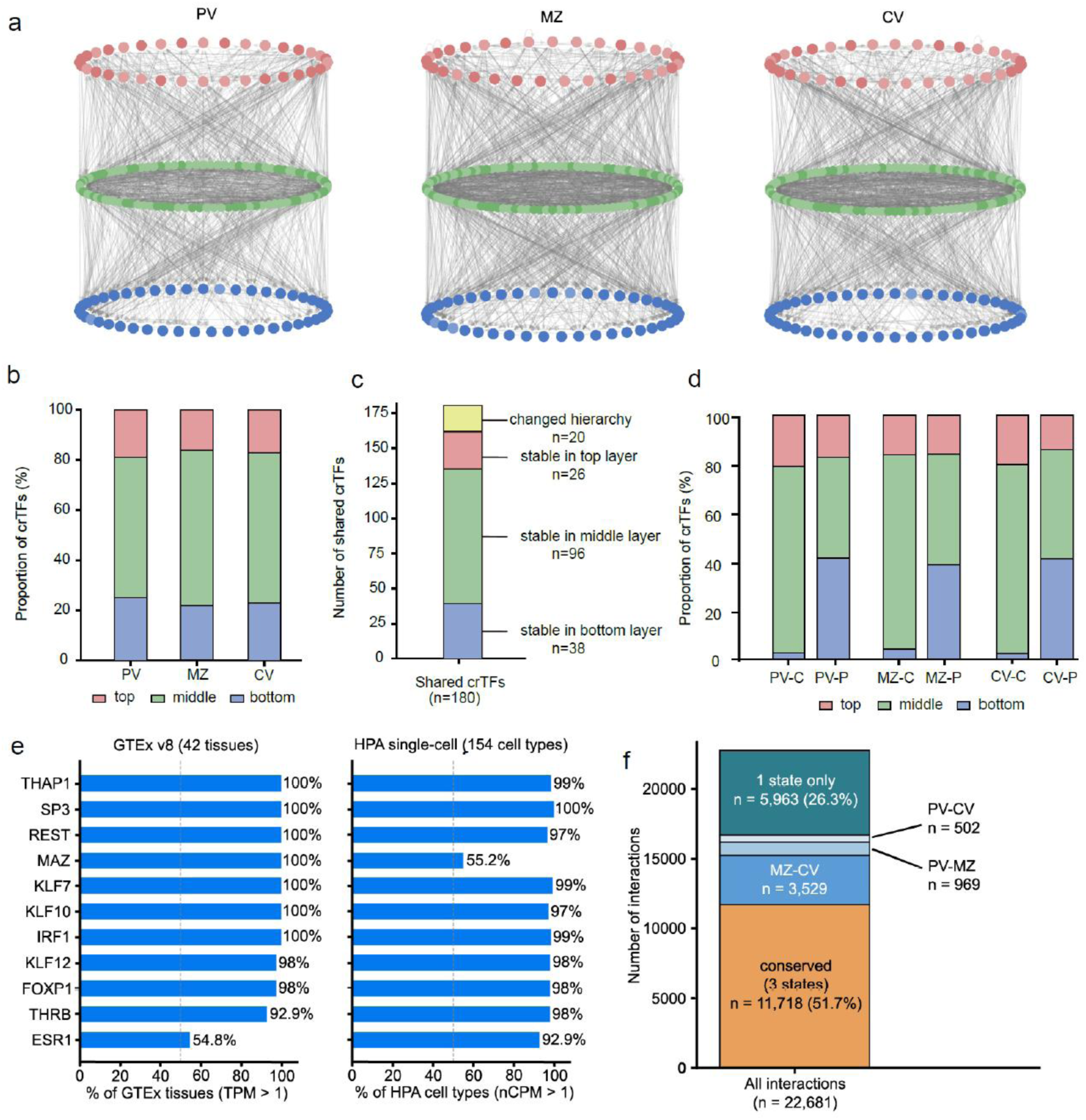
Comparative topological analysis identifies constitutive and state-specific components of the hepatocyte CRN. **a** Hierarchical visualization of CRNs across the three zones; for visual clarity, 10% of edges were randomly sampled. crTFs are color-coded as upstream (red), mediators (green), or effectors (blue). Among them, the light-colored nodes represent crTFs located in the core of the network. **b** Proportion of crTFs assigned to each hierarchical level across hepatocyte states. **c** Stability of hierarchical assignments for the 180 shared crTFs across all states. **d** Distribution of hierarchical levels within the network core (-C) and periphery (-P) partitions for each state. **e** Expression breadth of upstream-core crTFs. Fraction of GTEx v8 tissues (n = 42; TPM > 1) and Human Protein Atlas single-cell types (n = 154; nCPM > 1) in which each upstream-core crTF is expressed. Dashed line, 50%. **f** Variability of core regulatory interactions across states. Of 22,681 total interactions across the three state CRNs, 11,718 (51.7%) are conserved in all three states; of those present in two states, the majority are shared between MZ and CV (3,529), with fewer between PV and MZ (969) and PV and CV (502); 5,963 (26.3%) are present in a single state.

By comparing these two organizational levels within the 143 invariant crTFs, we found that the network core is a highly connected hub of upstream and mediator regulators, while the downstream effectors are largely positioned in the periphery (Fig. 5d). The upstream-core crTFs (namely, those that belong to both the structural core and the upstream hierarchical level) thus sit at the most prominent position of this architecture, providing regulatory signals to the rest of the network.

### Broadly expressed regulators form the upstream level of the invariant sub-network

Included among the 143 invariant crTFs are the established master regulators of hepatocyte identity, such as HNF4A, HNF1A, and FOXA2 (Dubois-Chevalier et al. 2023; Bravo González-Blas et al. 2024; Joo et al. 2019). Unexpectedly, within the CRN architecture, these master regulators were not found in the most upstream level, but rather at the mid-level within the network core (Fig. 5d and Supplemental Table S9) where they are involved in extensive interconnections with other crTFs. This central hub positioning is analogous to that of OCT4 and NANOG in embryonic stem cells, which are also considered central regulators (Boyer et al. 2005; Young 2011). This positioning suggests that their functioning as master regulators may actually be owing to such extensive interconnectedness within the network.

By contrast, the most upstream positions within the CRN architecture are occupied by a distinct subset that are located both at the network core as well as the upstream-level position (Fig. 5d and Supplemental Table S9). Strikingly, these upstream crTFs are not specifically expressed in hepatocytes. We examined their expression breadth using GTEx bulk RNA-seq across 42 human tissues and the Human Protein Atlas single-cell dataset across 154 cell types (Fig. 5e). We find 10 of 11 upstream-core crTFs are expressed in more than 92% of GTEx tissues (ESR1 the exception at 55%) and more than 92% of HPA single-cell types (MAZ the exception at 55%). By comparison, the master regulators HNF4A, HNF1A, and FOXA2 are expressed in only 19%, 17%, and 19% of GTEx tissues, respectively. Thus, the most upstream positions of the invariant sub-network are occupied by broadly expressed transcription factors, while tissue-specific regulators reside at mid-level integrative hubs.

Interestingly, these broadly expressed upstream-core factors are also implicated in liver diseases (Supplemental Table S10). For example, THRB is downregulated in hepatocellular carcinoma (HCC), and its activation has been shown to restore core TF expression in dedifferentiated hepatocytes (Ma et al. 2022). REST loss leads to de-repression of alternative lineage programs and is decreased in cholangiocarcinoma (Chong et al. 1995; Yu et al. 2018). Additional upstream-core factors, including KLF10, SP3, MAZ, and ESR1, are similarly implicated in HCC (Qiu et al. 2017; Pan et al. 2024; Luo et al. 2016).

### Variable interactions, composition and positioning define cell-state identity

Beyond the conserved features that define the hepatocyte cell type, each state CRN also clearly exhibits distinct features. At the interaction level, ∼27% of interactions among the 180 shared crTFs are reconfigured between states. When interactions involving state-specific crTFs are also considered, the extent of variability increases substantially: of the 22,681 total interactions across all three state CRNs, 10,963 (48.3%) are present in only one or two states (Fig. 5f; Supplemental Table S5). In pairwise terms, 31 to 38% of interactions differ between any two states (mean 35.5%; Supplemental Table S5). Among the interactions present in two states, the majority are shared between MZ and CV (3,529), with fewer between PV and MZ (969) and PV and CV (502), consistent with the spatial zonation gradient of the liver lobule. Notably, this rewiring is not concentrated on a subset of regulators but is distributed throughout the network: all 180 shared crTFs participate in at least one rewired interaction, and even the 10% most-rewired crTFs account for only ∼16% of rewired interactions (Supplemental Table S5).

At the compositional level, 39 crTFs are not shared across all three states, including 19 unique to a single zone and 20 present in only two states (Supplemental Table S6), with TCF7L1 specific to periportal hepatocytes and TBX3 and PPARG specific to pericentral hepatocytes (Bravo González-Blas et al. 2024; Cordoba-Chacon 2020). Additionally, 20 of the 180 shared crTFs shifted between core and periphery (Supplemental Table S7) and 20 changed hierarchical level (Supplemental Table S8) across states, including SREBF1, which resides in the core in CV but shifts to the periphery in PV. Thus, state-specific features occur at all three levels of network organization: composition, interactions, and positioning. When compared pairwise between states, these changes involve 23 to 29 crTFs, 20 to 29 positional shifts, and 30 to 40% of interactions being reconfigured (Jaccard distance). Notably, state specialization does not involve additional master regulators but rather a small set of largely zone-specific regulators. Thus, the compositional changes are modular, whereas the interaction rewiring is integrated across the network.

Importantly, these state-specific factors do not act independently of the invariant sub-network, as their binding motifs are found within the super-enhancer regions of identity-defining factors such as HNF4A and FOXA2 (Saint-André et al. 2016). Thus, state-specific factors act on the invariant sub-network rather than operating independently of it.

### The cell-type-level hepatocyte CRN resolves into invariant and state-dependent components

Finally, we asked how these state-level CRNs relate to the hepatocyte CRN previously defined at the level of the cell type (Dubois-Chevalier et al. 2023). That network comprises a core circuitry of 13 transcription factors, defined as a consensus of hepatocyte-identity factors across independent studies and confirmed by their co-recruitment at each other’s promoters in bulk mouse liver, together with a computationally derived extended network of 26 factors (Dubois-Chevalier et al. 2023). Despite the marked differences in network construction, 11 of the 13 core-circuitry factors are members of the invariant sub-network identified here, the two exceptions being HNF1B and NR0B2 (Supplemental Table S11). By contrast, of the 26 extended-network factors, over half are not invariant crTFs, including 7 not identified as crTFs in any state, 3 restricted to two of the three states, and 4 that, although shared across states, shift position (Supplemental Table S11). Thus, the experimentally established core circuitry was found to be part of the core, whereas the extended network is unreliable, indicating the limitations of the previous type-level characterization.

## Discussion

Previous models of CRNs have generally been derived from bulk tissue or cultured cell lines, which average over the multiple states present within a tissue. In this study, we developed LCM-Spatial-ATAC-seq, which achieves high-quality data from spatially defined cell populations within an intact tissue. Applying this approach to the mouse liver lobule, we generated state-resolved chromatin accessibility maps and, in combination with transcriptomic and chromatin-contact data, core regulatory networks for three hepatocyte states. To our knowledge, this is the first analysis of CRNs at state-level resolution.

Most notably, by comparing the different state-level CRNs, we could determine what is shared among all states, and is thus defining of a cell type, and what is different, and thus the characteristics of the states within this type. First and foremost, each state exhibits a very similar core-periphery and three-level hierarchy in their CRN. Thus, this appears to be a basic feature of these, perhaps all, cells. Beyond this, we found that cell-type identity is associated with three basic features: a shared composition of 143 crTFs present in all states, a conserved set of interactions (7,498) among these shared crTFs, and an invariant positioning within the core-periphery and hierarchical architecture. The cell-state CRNs have a dual character, consisting of this invariant sub-network together with state-specific differences in each of these features. To switch from one state to another would involve changing 20 to 30 crTFs, shifting the positions of 20 to 30 shared crTFs, and rewiring ∼36% of the interactions. To date, what distinguishes between cell states has remained somewhat enigmatic. Our results provide one quantitative description of this distinction. These state-level CRNs also resolve what the previous hepatocyte cell-type-level network (Dubois-Chevalier et al. 2023) actually represents: its experimentally established core circuitry corresponds to part of the invariant sub-network, whereas its extended network includes TFs that are invariant, state-dependent, and not identified in any state. Whether the type-level CRNs defined in other tissues similarly mix cell-type and state-dependent components remains an open question.

It may be illustrative to compare these findings with what was observed in the landmark paper by Saint-André et al., who identified CRNs in 75 different human cell types (Saint-André et al. 2016). Although their criteria for core TF membership differed from ours, and they did not analyze interactions or network structure, it is nonetheless intriguing to note that they observed a median fractional change of 92% in crTF composition between cell types (pairwise), in contrast to the 10 to 15% observed here between states. While recognizing that these differences may be owing to the notable differences in the analysis between the studies, it is tempting to speculate that this contrast reflects fundamental biological constraints on the extent of change a CRN can undergo and still remain a given cell type: small changes produce different states while larger changes produce a completely different type. In this light, it may be that cell types differ mainly in which crTFs are present, whereas cell states differ mainly in how a largely shared set of crTFs is wired.

However, we recognize that the distinction between states (and possibly types) is not solely quantitative, as our work has shown that different regulators have different roles in the CRN. In fact, one unexpected finding in our analysis is that the previously identified “master regulators” of cell identity are not at the most upstream positions in the regulatory hierarchy (Krauss and Kyba 2025). In particular, established hepatocyte master regulators, such as HNF4A, HNF1A, and FOXA2, reside at mid-level hub positions within the core, where they are extensively interconnected with other crTFs. Their master regulatory function may therefore derive not from being upstream of other regulators, but from their central, integrative position in the network, where both upstream and downstream signals are coordinated.

By contrast, the most upstream positions are occupied by broadly expressed factors. At first thought, this result is puzzling. One possible explanation is that only broadly expressed factors can suitably occupy this architectural position in the invariant sub-network. That is, broad expression indicates that these factors regulate, at least, generic pathways that many cell types need. By localizing to the most upstream positions, these factors are not embedded in many sub-networks within the CRN, especially compared to the middle-level regulators, leaving them isolated from transient changes in any cell-type-specific function that may be required. For example, it is known that HNF4A exhibits circadian-dependent activity (chromatin binding and isoform expression) (Qu et al. 2021; Deans et al. 2023). If this factor were located at the most upstream position, these oscillations in activity might jeopardize the entire homeostatic stability of the cell. Instead, by localizing to the middle, the effects of temporary changes in HNF4A expression can be local, with changes in the CRN itself mitigated by other members of its highly integrated sub-networks. Thus, the most upstream positions might be effectively reserved for regulators whose activity is critical to the cell at all times and in all states, impervious to cell-type-specific demands. As this would also be necessary for other cell types, this property could also explain the broad expression of these factors. Consistent with this view, independent work has shown that broadly expressed transcription factors are indeed strong stabilizers of cell identity (Missinato et al. 2023).

In addition, we should note that, while previous studies of CRNs focused on cell-type-specific TFs, broadly expressed TFs were, in fact, observed in the networks in at least some of this earlier work. In particular, in the aforementioned study by Saint-André et al., broadly expressed TFs were present in their identified networks (see Supplemental Table S3 in Saint-André et al. (2016)), although the authors considered these as auxiliary to the core CRNs of the cells (Saint-André et al. 2016). Similarly, in an extended network model of hepatocyte identity, Dubois-Chevalier et al. found that the core circuitry is complemented by additional TFs, the majority of which are not hepatocyte-specific, and that activation of two such factors can restore core TF expression in dedifferentiated hepatocytes (Dubois-Chevalier et al. 2023). Notably, 8 of 11 upstream-core crTFs described in our work here are also included in the Saint-André networks (Supplemental Table S11). Thus, we anticipate that broadly expressed TFs are a common feature of cell-type-defining CRNs more generally, but rather than being auxiliary, they actually play essential, upstream roles.

The upstream roles of these broadly expressed factors are reminiscent of what has been observed in other regulatory networks (Jothi et al. 2009; Yu and Gerstein 2006). At the whole-cell level, regulatory networks are organized hierarchically, with factors of more general function at the top (Gerstein et al. 2012). Developmental gene regulatory networks are organized similarly, with conserved sub-circuits upstream of more flexible differentiation programs (Erwin and Davidson 2009). By contrast, to date, the CRN has been treated as a set of mutually cross-regulating regulators, without a hierarchy (Almeida et al. 2021; Chubb et al. 2021). Our results may reconcile these views: the CRN is itself hierarchical, with broadly expressed, general-function factors at its top and the tissue-specific master regulators at mid-level hub positions. How this core hierarchy is embedded within the cell-wide regulatory network remains a fascinating question for future work.

The essential, upstream roles of these broadly expressed factors may also help explain why traditional cell reprogramming efforts, which have focused on the cell-type-specific factors, often yield unstable or functionally immature states (Du et al. 2014; Wang et al. 2021). Although broadly expressed, these upstream regulators are not necessarily part of the CRN of the starting cells, since CRN membership requires more than expression alone. Hence, switching the master regulators does not by itself establish this upstream layer of the target cell type. These upstream regulators may be essential stabilizers of the architecture needed for robust cell-type functionality. For example, it is known that the upstream activator THRB recruits the pBAF complex to remodel chromatin, enabling downstream hubs such as HNF4A to occupy their sites (Ma et al. 2022). Similarly, the broadly expressed factor REST has been found to actively repress alternative lineage programs (Chong et al. 1995). These activities, with upstream factors opening chromatin and protecting against changes in lineage, may provide important stabilizing functions that are needed to establish and maintain normal hepatocyte function.

Finally, we note that the structural stability of the 143 invariant crTFs suggests that this sub-network may be a critical site of disruption in disease. In hepatocellular carcinoma and advanced cirrhosis, cells often undergo dedifferentiation (Haep et al. 2021; Song et al. 2022), which could reflect stepwise disruption of this hierarchy. The disease associations we identified among these upstream regulators (Supplemental Table S10) are consistent with this view. Thus, just as the absence of these factors within the CRN may explain the instability of reprogrammed cells, the loss of these factors from normal hepatocytes could explain the devastating effects on the basic identity of the cell itself. Although these factors are more challenging to target owing to their broad expression, re-establishing their homeostatic expression may nonetheless be critical for effective therapy.

## Methods

### Preparation of the mouse fixed tissue samples

All animal procedures were approved by the Laboratory Animal Center of Shanghai Jiao Tong: University (approval. number: A2025020). and performed in accordance with institutional guidelines. The study is reported in accordance with the ARRIVE-guidelines. Eight- to ten-week-old male C57BL/6 mice (Jie Si Jie Laboratory Animals, Shanghai, China) were used. Mice were fed ad libitum and kept in a reverse light–dark cycle. Mice were euthanized by cervical dislocation at 6 h after lights off. The liver and kidney were freshly dissected, cleaned in ice-cold phosphate-buffered saline (PBS) solution, and then cut into small blocks. Following optimization, liver and kidney blocks were fixed in 1% PFA (17 °C) for 2.5 and 4 h, respectively. Fixation was quenched with 0.125 M glycine for 15 min at room temperature and then for 15 min at 4 °C. Finally, the samples were maintained on ice for subsequent sectioning.

### *In situ* ATAC-seq performed in tissue sections

Tissue blocks were put into ice-cold PBS and sectioned at a thickness of 10 µm using a vibratome (speed: 0.30 mm/s, amplitude: 0.75). ATAC-seq was performed following a published method (Corces et al. 2017) with modifications for free-floating tissue sections. For the optimized procedure, the sections were incubated in 0.125 M glycine for 5 min, washed twice with 1 x PBS, and cleaned in ddH2O. The sections were permeabilized with 1 mL of lysis buffer (10 mM Tris-HCl, pH 7.4, 10 mM NaCl, 3 mM MgCl2, 0.1% Tween-20, 0.1% NP-40, and 0.01% digitonin) for 20 min on ice, followed by a 10-min wash in 1 mL of wash buffer (10 mM Tris-HCl, pH 7.4, 10 mM NaCl, 3 mM MgCl2, and 0.1% Tween-20). To standardize the Tn5-to-DNA stoichiometry, we first calibrated the relationship between tissue area and genomic DNA content. Based on this calibration, the tissue area used in each reaction was converted into an estimated number of cells, and the transposition reaction was scaled proportionally to maintain a constant Tn5-to-DNA ratio. A 50-µL transposition reaction was used for tissue sections estimated to contain approximately 5 × 10⁴ cells, and larger tissue inputs were processed by increasing all reaction components proportionally. The 50-µL reaction consisted of 25 µL 2× TD Buffer, 9 µL 1× PBS, 0.5 µL 10% Tween-20, 0.5 µL 1% digitonin, 10 µL 5× TTBL and 5 µL TTE Mix. The reaction was incubated at 37 °C for 60 min with gentle shaking at 300 rpm on a ThermoMixer. After removing the transposition mix, 500 µl 40 mM EDTA was added for incubation at room temperature for 5 min to stop transposition. Finally, the sections were washed with 500 µl of 1× NEBuffer 3.1 (B6003S, NEB) for 5 min at room temperature and stored at -80 °C until further processing.

### Immunofluorescence Staining

Sections were washed three times with 1× PBS for 5 min each and then blocked with 5% BSA (A7030, Sigma-Aldrich) for 1h at room temperature. For liver tissue staining, sections were then incubated overnight at 4 °C with mouse anti-Glutamine Synthetase (GS) mAb (1:200; GB121177, Servicebio) and rabbit anti-E-Cadherin (24E10) mAb (1:100; 3195T, Cell Signaling Technology). GS and E-cadherin staining were used to identify pericentral and periportal hepatocytes, respectively, whereas mid-zonal hepatocytes were defined as the unstained region located between the GS-positive and E-cadherin-positive regions. For kidney tissue staining, sections were incubated overnight at 4 °C with rabbit anti-EpCAM mAb antibody (1:200; 42515S, Cell Signaling Technology) to identify epithelial cells. Following three 5-min washes with PBST (1× PBS containing 0.1% Tween-20), sections were incubated for 2 h at room temperature with secondary antibodies in the dark with the corresponding secondary antibodies: Alexa Fluor Plus 488-conjugated donkey anti-mouse IgG (1:200; A32766, Invitrogen) for GS staining and Alexa Fluor 568-conjugated donkey anti-rabbit IgG (1:200; A10042, Invitrogen) for E-cadherin or EpCAM staining. After three final 5-min washes with PBST, sections were immersed in ddH2O and then processed for subsequent microdissection.

### Laser Capture Microdissection

Following immunofluorescence staining, the sections were placed on a polyethylene naphthalate (PEN)-membrane-covered slide (415190-9041-000, Carl Zeiss). A PEN-membrane-covered slide was treated with UV irradiation for 30 min prior to mounting the sections. Desired cells were manually selected on the control screen based on their fluorescence signal using the Zeiss PALM Micro Beam LCM system (Zeiss Microimaging, Munich, Germany). Before microdissection, excess solution was removed with a pipette from the slide and the sections were allowed to dry fully on the PALM stage. The materials dissected with the UV laser were ejected into 0.2 ml adhesive cap tubes (415190-9191-000, Carl Zeiss). We continued this process, collecting material one cell at a time until sufficient materials were accumulated. To prevent contamination, all LCM steps were performed within a dedicated plexiglass chamber. We also inspected the adhesive cap of the collection tubes under phase-contrast microscopy to confirm the absence of debris or foreign particulates, and subsequently placed the tube at 4 °C to perform the subsequent experimental steps.

### DNA Extraction and ATAC-seq Library Preparation

The collected samples were incubated in a reverse crosslinking solution (50mM Tris-HCl, pH 8, 1mM EDTA, 0.4 mg/mL proteinase K, 0.1% SDS, and 200 mM NaCl) at 65 °C overnight. The DNA was purified using 1.8× Agencourt AMPure XP beads (A63881, Beckman) according to the manufacturer’s instructions and resuspended in 10 mM Tris-HCl, pH 8.0. The amount of LCM-captured material was estimated from the DNA recovered after reverse cross-linking and purification, using ∼6 pg DNA per mouse cell as a reference. Subsequently, libraries were then generated by PCR amplification using TruePrep DNA Library Prep Kit V2 (TD501, Vazyme). The concentration and size distribution of the resulting libraries were quantified using an Agilent Bioanalyzer High Sensitivity DNA Chip. Finally, all libraries were sequenced on the Illumina NovaSeq 6000 platform using paired-end 150-bp reads.

### ATAC-seq data analysis

Adapter-contaminated and low-quality bases were removed from the raw sequencing reads using trim_galore (v0.6.6), using the parameter “-q 20 --stringency 1 --length 20 -e 0. --paired”. After trimming, reads were mapped to mm10 reference genome with Bowtie2 (v2.3.4.3) (Langmead and Salzberg 2012), the parameters “-q - X 1000 --very-sensitive -k 1 --no-mixed -p 8” was used. Using Sambamba (v0.6.6) (Tarasov et al. 2015), we filtered the uniquely mapped reads, measured the mapping ratio. Reads aligned to mtDNA, reads with low mapping quality (MAPQ <30), and ENCODE blacklist regions were removed using SAMtools. Sambamba was then used to remove duplicate reads. ATAC libraries from individual sorting rounds per sample were sequenced and pooled to generate each biological replicate. For peak calling, deduplicated nuclear genome-mapped reads from the two biological replicates of each condition were merged. To ensure comparable sequencing depth across conditions, each merged BAM file was down-sampled to the lowest number of nuclear genome-mapped reads among all conditions. Then, Genrich software was used to call peaks with the ATAC-seq mode (-r -t (input.bam) -j -a 100) (https://github.com/jsh58/Genrich).

### Calculation of ATAC-seq QC metrics

ATAC-seq data quality was evaluated using three quality-control metrics: transcription start site (TSS) enrichment, insert size distribution and fraction of reads in peaks (FRiP), following the ENCODE ATAC-seq Data Standards and Processing Pipeline (https://www.encodeproject.org/atac-seq/). TSS enrichment scores were calculated using TSSenrich v1.4.6. Insert size distributions were assessed using ATACseqQC v1.10.4 in R, and FRiP scores were calculated as the fraction of mapped reads overlapping called peak regions. Samples with TSS enrichment scores >10 were considered acceptable. FRiP scores >0.3 were considered preferred, whereas values >0.2 were considered acceptable. Insert size distribution was inspected to confirm the expected ATAC-seq fragment size periodicity.

### Identification of highly accessible peaks

Early studies suggested that the distribution of mapped ATAC-seq reads follows a Poisson distribution parameterized by λ = (total read counts × read length) / genome length. To address potential biases, we incorporated a control dataset of Tn5-accessible regions in nucleosome-free genomic DNA, enabling systematic correction of Tn5 sequence bias. This established a statistically validated Poisson model for the experimental data.

To quantify chromatin accessibility peaks, we computed the cumulative probability for each peak under this Poisson model. Specifically, for a peak containing k normalized reads within a genomic window of size L, the cumulative probability p was calculated as:

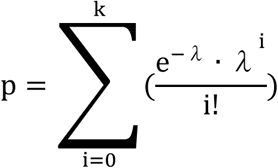

where λ = (total normalized reads counts × reads length) /effective length(L), representing the expected reads count under the null model of uniform accessibility. Peaks exhibiting p > 0.9 were classified as high-confidence open chromatin regions, indicating ≥ 90% probability that the observed read density exceeds random background distribution.

### Identification of hepatocyte super enhancers and assignment of state activity

Super-enhancers (SEs) were identified using the Rank Ordering of Super-Enhancers (ROSE) algorithm (Whyte et al. 2013). To avoid potential confounding signals from active promoters, regions within 1 kilobase (kb) of transcription start sites (TSSs) were excluded (parameter -t 1000). Adjacent H3K27ac ChIP-seq peaks were stitched together within a 12.5 kb range (parameter -s 12500). Subsequently, Stitched enhancer regions were then ranked according to their H3K27ac signal intensity, and regions above the ROSE-defined inflection-point cutoff were classified as SEs. The published H3K27ac ChIP-seq data for hepatocytes is available in GSM3135566. Because the H3K27ac ChIP-seq data were generated from hepatocytes comprising all zonated states, SEs were first identified at the hepatocyte level as described above and then assigned to individual hepatocyte states using state-specific chromatin accessibility. Specifically, an SE was considered active in a given state if it overlapped, by at least 1 bp, with at least one highly accessible ATAC-seq peak in that state. These state-active SEs were used for downstream assignment to expressed TF genes and construction of hepatocyte state-specific CRNs.

### RNA-seq data analysis

Mouse liver RNA-seq data were downloaded from the NCBI Sequence Read Archive (SRA) with the BioProject accession code PRJNA556572 and analyzed using the zUMIs pipeline. Samples from different liver regions were distinguished based on barcodes information provided by Ben-Moshe et al. (2019). Zonation region 1 and 2 were classified as CV hepatocytes, zonation region 4 and 5 were classified as MZ hepatocytes, and zonation region 7 and 8 were classified as PV hepatocytes. Reads were aligned to Mus_musculus.GRCm38.96.gtf downloaded from https://www.ensembl.org/index.html using STAR (v2.7.11a). Exonic unique molecular identifier read counts were calculated with default parameters. TF genes were annotated using the mouse TF catalogue from AnimalTFDB 4.0. For each hepatocyte state, TF genes with CPM > 1 were considered expressed in that state.

### Hi-C data analysis

Hi-C reads were iteratively mapped to the mm10 genome using Bowtie2 (v2.1.0) with a step size of 25 bp. Reads mapped to multiple genomic locations or with low mapping quality, defined as MAPQ < 30, were removed. PCR duplicates were further removed. Read pairs identified as same fragment (i.e., pairs mapped to the same restriction fragment), nonspecific ligations (fragments with sizes above 600 bp), dangling (inward-facing read pairs less than 1 kbp apart), and self-ligations (outward-facing read pairs less than 25 kbp apart) were excluded. Valid read pairs were retained for each sample Juicer tools (v1.19.02) were used to create the .hic file and perform KR normalization. Significant chromatin contacts were identified using FitHiC2 (Kaul et al. 2020). Bins with sparse coverage, defined as bins contacting fewer than 3% of other bins, were excluded from the analysis. Based on our sequencing depth, a 20-kb bin size was selected to ensure robust contact detection. Significant contacts were defined as FitHiC2 interactions with q value < 0.05. SE-associated contacts were identified by intersecting significant contact loci with SE regions. The published liver Hi-C data are available in NCBI under the accession numbers PRJNA1187272. For each hepatocyte state, an SE was assigned to a TF gene when one 20-kb bin of a significant contact overlapped a state-active SE and the paired 20kb-bin contained the TSS of a TF gene expressed in the same state. TF genes linked to at least one state-active SE through such significant contacts were defined as state-specific SE-associated TFs.

### Construction of hepatocyte state-specific core regulatory networks

For each hepatocyte state, sequences of state-highly accessible ATAC-seq peaks located within state-active SEs associated with expressed TF genes were extracted and scanned for TF-binding motifs using FIMO v5.0.2. Position-weight matrices were obtained from JASPAR 2024, and motif matches were retained at *p* < 1 × 10⁻⁵. Motif scanning was restricted to state-specific SE-associated TFs with annotated motifs. A directed edge from TF A to TF B was assigned when a motif for TF A was identified within a state-highly accessible peak located in a state-active SE associated with TF B. Edges were assembled into a state-specific TF interaction network, and TFs participating in at least one motif-supported interaction were defined as crTFs.

### Neighborhood Similarity Across Multiple Networks

For the three directed networks CV, PV, and MZ, we focus on their common node set and quantify how consistently each node maintains its neighborhood across the networks. For a node u, let N_CV_(u), N_PV_(u), and N_MZ_(u) denote its neighbor sets in the three networks, respectively, we define the neighborhood similarity of u using the Jaccard similarity index, a standard measure of set overlap:

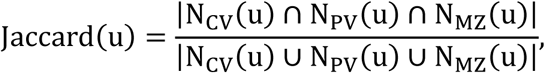

where | ⋅ | denotes taking the size of a set. Jaccard(u) captures the extent to which the neighborhoods of u are stable across different networks. Higher value indicate that u’ s connectivity pattern is stable, conserved, or robust, suggesting that it may play a structurally important or functionally conserved role across different networks. Focusing on the regulatory property, we computed this score for each common node using the union of its incoming and outgoing neighbors in CV, PV and MZ, and then ranked the nodes to identify those with the highest neighbor similarity. After computing this score for every common node, we remove nodes whose similarity is zero, as these cases arise when a node has no shared out-neighbors across CV, PV, and MZ and therefore carry limited structural information.

### Null Models in Network Analysis

In our analysis, we considered two null models: Erdős–Rényi (ER) (Erdős and Rényi 1959) and directed binary configuration model (DBCM) (Squartini and Garlaschelli 2011). ER does not control the degree while DBCM accounts for the degree heterogeneity. For the ER null, we preserve the non-self-loop edge density and the number of self-loops separately. We first compute the density *p_e_* from the observed non-self-loop edges, and generate a directed ER network on the same node set by independently sampling each possible non-self-loop edge with probability *p_e_*. We then randomly insert the same number of self-loops as in the original network. To generate a DBCM randomized network, we solve for *x_i_* and *y_i_* so that the expected in- and out-degrees match the observed network. Each ordered non-self-loop edge is then sampled independently with probability *p_ij_* = *x_i_y_j_*/(1 + *x_i_y_j_*).

### Community and Core–Periphery Detection in Networks PV, MZ and CV

To search for community structure, we applied five complementary methods. Directed Louvain (Dugué and Perez 2015) and OSLOM (Lancichinetti et al. 2011) evaluate community structure relative to degree-based null models. Infomap (Rosvall and Bergstrom 2008) and the Girvan-Newman (GN) algorithm (Girvan and Newman 2002) do not explicitly incorporate degree correction in generating their partitions. For the hierarchical stochastic block model (hSBM) (Peixoto 2014), we fitted both degree-corrected and non-degree-corrected variants and selected the fit with the smallest description length.

For each method, we scanned its main tuning parameter. For Louvain, we varied the resolution parameter γ over [0.5, 0.75, 1.0, 1.5, 2.0]. For OSLOM, we varied the module significance threshold t over [0.05, 0.10, 0.20, 0.50] and the coverage parameter cp over [0.1, 0.5, 0.9] for first-level detection. For Infomap, we varied the Markov time over [0.5, 0.75, 0.9, 1.0, 1.5, 2.0]. For GN, we cut the dendrogram at levels [2, 5, 10, 20, 50, 100]. For hSBM, instead of specifying a resolution parameter, we allowed MDL to select the optimal models. None of the methods yielded robust non-trivial community structure in any network state over the parameter ranges considered. For Directed Louvain, the optimized modularity never exceeded approximately *Q* ∼ 0.05, whereas the modularity of the GN partitions remained below 0.01. Infomap returned either a single module or one module accompanied only by singleton modules, and OSLOM identified no statistically significant modules. For hSBM, the minimum-description-length fit was the trivial one-block model in every state. Taken together, both degree-adjusted and unadjusted approaches provided no robust evidence of non-trivial community structure in the CRNs within the methods and parameter ranges considered.

For core-periphery detection, we applied Lip’s method to identify potential core-periphery structures, which is an efficient and parameter-free core-periphery detection method. Since it is designed for undirected graphs, we symmetrized the adjacency matrix of each network and applied Lip’s method to obtain the detection results.

The detected core-periphery structure is highly significant against the ER null in all three states (*p* ∼ 0.005), and is not significant against the DBCM null (We use the DBCM rather than the Chung-Lu model of Yanchenko et al. (2025), since our CRNs are dense enough that Chung-Lu probabilities exceed unity; the DBCM is its exact counterpart). The core-periphery structure of the CRNs can therefore be understood as an exogenous core-periphery structure, consistent with Yanchenko et al. (2025).

### Hierarchical analysis of hepatocyte state-specific CRNs

All crTFs in each state-specific CRN were classified into three hierarchical levels based on network topology using a previously described hierarchy height metric (Gerstein et al. 2012; Song et al. 2016). For each crTF, out-degree (O), defined as the number of downstream crTFs predicted to be regulated by that crTF, and in-degree (I), defined as the number of upstream crTFs predicted to regulate that crTF, were calculated from the directed CRN. A normalized hierarchy height was then defined as ℎ = (*O* − *I*)/(*O* + *I*). crTFs were classified as top-level regulators when 0.33 < ℎ ≤ 1, bottom-level regulators when −1 ≤ ℎ < −0.33, and middle-level regulators when −0.33 ≤ ℎ ≤ 0.33. The significance of the distribution of h metric of TFs, was calculated from the distribution of h in 1000 random networks (KS tests). For each empirical network, 1000 randomized directed networks were generated using the ER model. The KS statistic was defined as the average KS distance between the empirical and randomized h distributions, and its empirical p-value was calculated against a null distribution of KS statistics obtained from random-network pair comparisons.

## Public Datasets

Publicly available datasets used in this study were obtained from GEO (https://www.ncbi.nlm.nih.gov/geo/), including unfixed HCT116 ATAC-seq data under accession GSE170407 and hepatocyte H3K27ac ChIP-seq data under accession GSM3135566, and from the NCBI Sequence Read Archive (https://www.ncbi.nlm.nih.gov/sra), including fresh frozen mouse liver and kidney bulk ATAC-seq data under accession SRP167062 and liver Hi-C data under BioProject accession PRJNA1187272. The GTEx bulk RNA-seq data and Human Protein Atlas single-cell data used for the expression-breadth analysis were obtained from the GTEx portal (https://gtexportal.org) and the Human Protein Atlas (https://www.proteinatlas.org).

## Software availability

The analysis scripts used in this study are available at GitHub: https://github.com/rzhou-dev/CRN-Spatial-Mapping.

## Data Access

The raw sequencing reads generated in this study are available in the repositories of the National Genomics Data Center (NGDC; https://ngdc.cncb.ac.cn), under BioProject accession HRA018756 and CRA043792. Processed ATAC-seq signal tracks can be visualized through the UCSC Genome Browser session at https://genome.ucsc.edu/s/Zhourui/mm10.

## Acknowledgments

We thank all members of the Shao Laboratory (Shanghai, China) for their help in critical comments during the execution of this study. We thank Genefund Biotech (Shanghai, China) for assistance with data analysis. We thank Tianchen Deng (Shanghai, China) for helpful discussions on computational analysis. This work was supported by grants from the National Key R&D Program of China (No. 2023YFE0108800), Shanghai Pilot Program for Basic Research, the National Natural Science Foundation of China (Grant Nos. 31971151, 81627801, 81972909, and 32370572), and the KC Wong Education Foundation (Hong Kong, China). Biomni (Phylo) was used to assist with manuscript text revision and preparation of the literature-based annotations and references in Supplemental Table S10. The authors take responsibility for the final content.

## Author Contributions

Rui Zhou: Investigation, Methodology, Formal analysis, Writing - original draft. Li Ge: Software, Formal analysis. Jing Li: Methodology. Zeqian Xu: Formal analysis. Min Chen: Formal analysis. Yan Guo: Methodology. Daniel M. Czajkowsky: Conceptualization, Formal analysis, Writing - original draft. Zhifeng Shao: Conceptualization, Methodology, Writing - original draft. All authors read and approved the final manuscript.

