## Supplemental Figures and Text for "An invariant network architecture with broadly expressed regulators distinguishes cell-type from cell-state identity *in situ*"

### Supplemental figures and legends

Supplemental Fig. S1. Additional quality assessment of fixed-section bulk and LCM-EpCAM ATAC-seq datasets.
Supplemental Fig. S2. Quality assessment of ATAC-seq datasets from PV, MZ, and CV hepatocytes.
Supplemental Fig. S3. State-resolved accessibility landscapes validate biological zonation.
Supplemental Fig. S4. Quality control and intermediate analyses for CRN construction.
Supplemental Fig. S5. Comparison of h distributions between empirical and randomized networks across CV, PV, and MZ

### Supplemental tables

Supplemental Table S1. Optimization and quality assessment of ATAC-seq libraries from fixed HCT116 cells.
Supplemental Table S2. Optimization and quality assessment of fixed mouse tissue section ATAC-seq.
Supplemental Table S3. Accessible peaks, fully accessible peaks, and active SEs in PV, MZ, and CV hepatocytes.
Supplemental Table S4. State-expressed TF genes and Hi-C-supported SE-associated TF genes in PV, MZ, and CV hepatocytes.
Supplemental Table S5. crTFs and CRN edges in PV, MZ, and CV hepatocytes.
Supplemental Table S6. Distribution and neighborhood similarity of crTFs across PV, MZ, and CV hepatocyte CRNs.
Supplemental Table S7. Core–periphery classification of crTFs across PV, MZ, and CV hepatocyte CRNs.
Supplemental Table S8. Hierarchical classification of crTFs in PV, MZ, and CV hepatocyte CRNs.
Supplemental Table S9. Conserved shared crTFs across PV, MZ, and CV hepatocytes.
Supplemental Table S10. Liver-disease associations of broadly expressed upstream-core crTFs.
Supplemental Table S11. Classification of published cell-type-level hepatocyte CRN factors in the state-level CRNs.

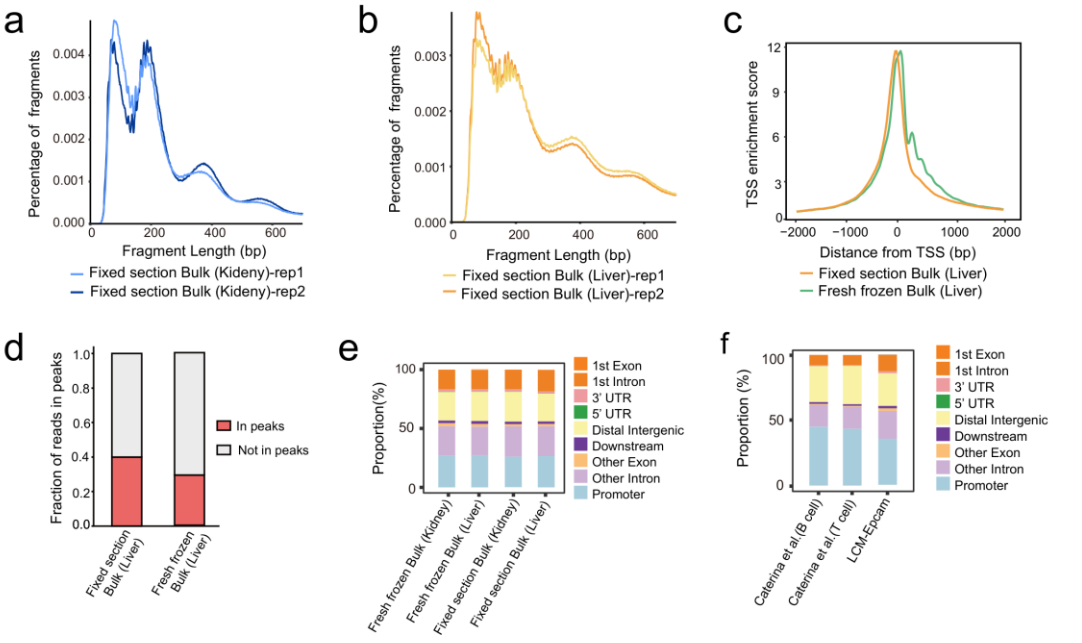


**Supplemental Fig. S1. Additional quality assessment of fixed-section bulk and LCM-EpCAM ATAC-seq datasets.**

**a,b** Fragment length distributions (FLDs) of fixed-section bulk ATAC-seq datasets generated from mouse kidney (**a**) and liver (**b**) tissues. Two biological replicates are shown for each tissue. **c** Comparison of transcription start site (TSS) enrichment profiles between merged fixed-section bulk ATAC-seq data from mouse liver tissue and published fresh-frozen bulk liver ATAC-seq data. **d** Fraction of reads in peaks (FRiP) scores for fixed-section bulk liver ATAC-seq data and published fresh-frozen bulk liver ATAC-seq data. **e** Genomic annotation of ATAC-seq peaks from fixed-section bulk and published fresh-frozen bulk ATAC-seq datasets from mouse kidney and liver tissues. **f** Genomic annotation of ATAC-seq peaks from LCM-EpCAM ATAC-seq data, fixed-section bulk kidney ATAC-seq data, and published LCM-captured B cell and T cell ATAC-seq datasets. FLD, fragment length distribution; TSS, transcription start site; FRiP, fraction of reads in peaks; LCM, laser capture microdissection.

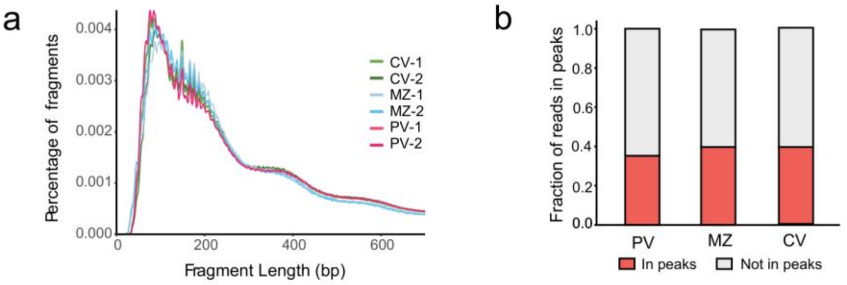


**Supplemental Fig. S2. Quality assessment of ATAC-seq datasets from PV, MZ, and CV hepatocytes.**

**a** Fragment length distribution (FLD) of ATAC-seq data from two biological replicates of PV, MZ, and CV hepatocytes. **b** Fraction of reads in peaks (FRiP) for two biological replicates of PV, MZ, and CV hepatocytes. Red indicates reads in peaks, and grey indicates reads not in peaks. PV, portal vein-associated hepatocytes; MZ, mid-zone hepatocytes; CV, central vein-associated hepatocytes.

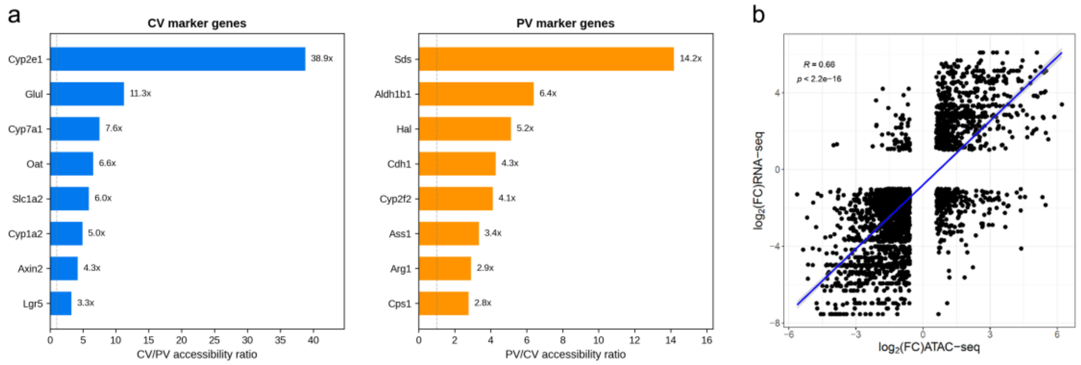


**Supplemental Fig. S3. State-resolved accessibility landscapes validate biological zonation.**

**a** Accessibility ratios at established zonation marker genes. Left: CV marker genes, shown as the CV/PV accessibility ratio; right: PV marker genes, shown as the PV/CV accessibility ratio. Cdh1 and Glul, the antigens used for PV and CV immunolabeling, respectively, show the expected zone-specific accessibility. Dashed line indicates equal accessibility (ratio = 1). **b** Correlation between changes in chromatin accessibility and gene expression (CV versus PV) among differentially expressed genes. log2 fold-change in promoter accessibility (x-axis) versus log2 fold-change in expression (y-axis); Pearson correlation coefficient and p-value are indicated.

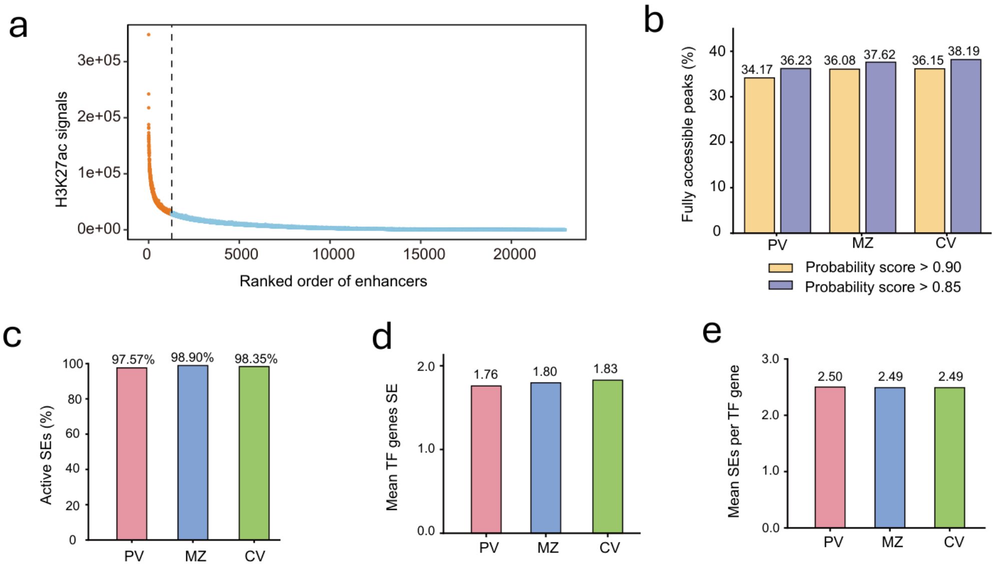


**Supplemental Fig. S4. Quality control and intermediate analyses for CRN construction.**

**a** ROSE rank-ordering plot of enhancers identified from published hepatocyte H3K27ac ChIP-seq data (Li et al. 2019). Enhancers were ranked by H3K27ac ChIP-seq signal intensity. Super-enhancers are shown in orange, and typical enhancers are shown in blue. The dashed vertical line indicates the cutoff used to define super-enhancers. **b** Fraction of highly accessible peaks identified in PV, MZ and CV hepatocytes using probability score cutoffs of >0.90 and >0.85. Peaks with probability scores >0.90 were used as the final set of highly accessible peaks. **c** Percentage of super-enhancers (SEs) identified as active in each hepatocyte zone. **d** Mean number of expressed transcription factor (TF) genes linked to each active SE in PV, MZ, and CV hepatocytes. **e** Mean number of active SEs linked to each expressed TF gene in PV, MZ, and CV hepatocytes. PV, periportal hepatocytes; MZ, mid-zone hepatocytes; CV, pericentral hepatocytes; ROSE, Rank Ordering of Super-Enhancers; SE, super-enhancer; TF, transcription factor.

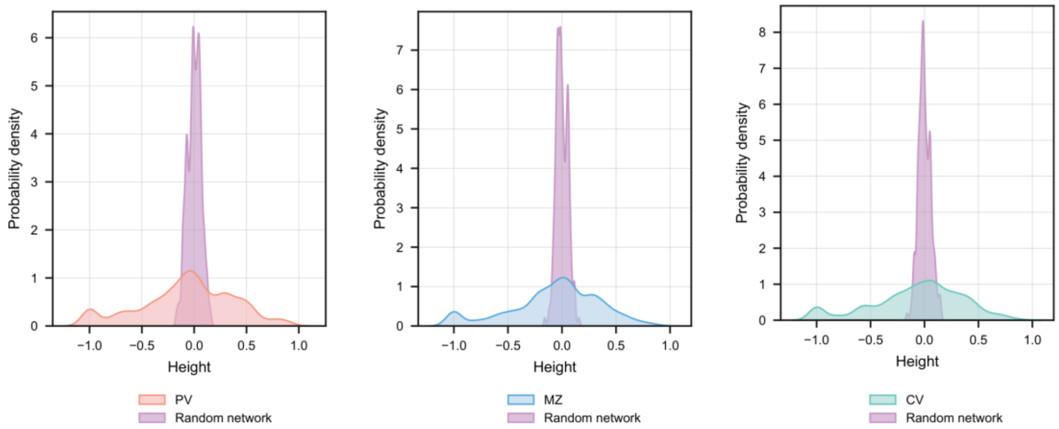


**Supplemental Fig. S5. Comparison of h distributions between empirical and randomized networks across CV, PV, and MZ**

Kernel density estimate of the h distribution in the empirical network and one randomized network for CV, PV, and MZ. One randomized network is shown for visualization, while significance was assessed using the KS statistic as a distribution-level distance measure based on 1000 randomized networks for each network. The empirical *p* value was <0.001 for all three networks.

**Supplemental Table S1. Optimization and quality assessment of ATAC-seq libraries from fixed HCT116 cells.**

Worksheet 1 summarizes the optimization of PFA fixation conditions and Tn5 tagmentation parameters in HCT116 cells, including fixation time, PFA concentration, Tn5 concentration, and Tn5 reaction time. The optimal condition was selected based on the highest TSS enrichment score. Worksheet 2 compares ATAC-seq quality metrics between optimally fixed HCT116 cells, unfixed HCT116 cells prepared in this study, and unfixed HCT116 ATAC-seq datasets obtained from ENCODE. Fixed HCT116 libraries were generated using the optimized condition identified in Worksheet 1, consisting of 0.1% PFA fixation for 5 min at room temperature followed by 1× Tn5 tagmentation for 60 min. All replicates were downsampled to 20 million valid paired-end fragments before calculating TSS enrichment scores and FRiP values. Values are shown for individual replicates. PFA, paraformaldehyde; Tn5, Tn5 transposase; TSS, transcription start site; FRiP, fraction of reads in peaks.

**Supplemental Table S2. Optimization and quality assessment of fixed mouse tissue section ATAC-seq.**

Worksheet 1 summarizes the optimization of fixation conditions for fixed mouse kidney tissue section ATAC-seq. Mouse kidney tissue blocks were fixed with 1% PFA at 17 °C for the indicated durations, followed by ATAC-seq on free-floating tissue sections using 1× Tn5 tagmentation for 60 min. Worksheet 2 compares ATAC-seq quality metrics between fixed section bulk kidney samples generated in this study and published fresh frozen bulk kidney datasets. Worksheet 3 summarizes the optimization of fixation and Tn5 tagmentation conditions for fixed mouse liver tissue section ATAC-seq. Mouse liver tissue blocks were fixed with 1% PFA at 17 °C under the indicated conditions, and an increased Tn5 concentration was tested to reduce the high proportion of mitochondrial reads observed in liver samples. Worksheet 4 compares ATAC-seq quality metrics between fixed section bulk liver samples generated in this study and published fresh-frozen bulk liver datasets. In the optimization worksheets, the optimal condition was selected based on TSS enrichment score and mitochondrial read fraction. In the quality comparison worksheets, all replicates were downsampled to 20 million valid paired-end fragments before calculating TSS enrichment scores and FRiP values. Values are shown for individual replicates where applicable. PFA, paraformaldehyde; Tn5, Tn5 transposase; TSS, transcription start site; FRiP, fraction of reads in peaks.

**Supplemental Table S3. Accessible peaks, fully accessible peaks, and active SEs in PV, MZ, and CV hepatocytes.**

Worksheet 1 lists accessible peaks identified in PV, MZ, and CV hepatocytes; each biological replicate was downsampled to 20 million valid paired-end fragments before pooling replicates from the same hepatocyte state for peak calling with Genrich in ATAC-seq mode. Worksheet 2 lists hepatocyte SEs identified from published H3K27ac ChIP-seq data using ROSE. Worksheet 3 lists highly accessible peaks identified in each hepatocyte state; each ATAC-seq peak was assigned a probability score based on the likelihood of observing its read counts under a randomized distribution, and peaks with probability scores >0.90 were defined as fully accessible. Worksheet 4 lists ROSE-defined SEs that were defined as active in a given hepatocyte state if they overlapped at least one fully accessible peak in that state.

Worksheet 5 lists fully accessible peaks located within active SEs, which were used for downstream motif analysis. Genomic coordinates are based on the mm10 mouse genome assembly. SE, super-enhancer; ROSE, Rank Ordering of Super-Enhancers; PV, portal vein-associated hepatocytes; MZ, mid-zone hepatocytes; CV, central vein-associated hepatocytes.

**Supplemental Table S4. State-expressed TF genes and Hi-C-supported SE-associated TF genes in PV, MZ, and CV hepatocytes.**

Worksheet 1 provides summary statistics for state-expressed TF genes and SE-associated TF genes, including the numbers of expressed TFs, unique SE–TF pairs, unique SE-associated TF genes, unique SEs linked to TF genes, and mean numbers of TF genes per SE and SEs per TF gene. Worksheet 2 lists mouse TF genes annotated using the AnimalTFDB v4.0 mouse TF annotation; TF genes with CPM >1 in a given hepatocyte state were defined as expressed in that state. Worksheets 3–5 list SE-associated TF genes in PV, MZ, and CV hepatocytes, respectively. SE-associated TF genes were identified by linking state-specific active SEs to TF genes expressed in the corresponding hepatocyte state using statistically significant chromatin contacts from mouse liver Hi-C maps with FitHi-C. Significant contacts were defined using a $q$-value cutoff of <0.05. CPM, counts per million; SE, super-enhancer; TF, transcription factor.

**Supplemental Table S5. crTFs and CRN edges in PV, MZ, and CV hepatocytes.**

Worksheet 1 provides summary statistics for crTFs and CRN edges in each hepatocyte state. Worksheets 2–4 list CRN edges in PV, MZ, and CV hepatocytes, respectively. Edges are defined as regulator–target TF pairs linked through active SEs containing fully accessible ATAC-seq peaks with matching regulator motifs. Worksheet 5 lists the crTFs identified in each hepatocyte state. crTF, core regulatory transcription factor; CRN, core regulatory network; SE, super-enhancer; TF, transcription factor.

**Supplemental Table S6. Distribution and neighborhood similarity of crTFs across PV, MZ, and CV hepatocyte CRNs.**

Worksheet 1 lists the presence of each crTF across the three hepatocyte states and classifies crTFs as shared across all three states, present in two states, or state-specific. Worksheet 2 lists neighborhood similarity scores for shared crTFs across PV, MZ, and CV CRNs. NSS, neighborhood similarity score.

**Supplemental Table S7. Core–periphery classification of crTFs across PV, MZ, and CV hepatocyte CRNs.**

The table lists the core or periphery assignment of each crTF in each state and compares the structural assignments of shared crTFs across PV, MZ, and CV.

**Supplemental Table S8. Hierarchical classification of crTFs in PV, MZ, and CV hepatocyte CRNs.**

crTFs were classified into top, middle, and bottom roles based on their in-degree and out-degree connectivity in each hepatocyte CRN. A hierarchy score was calculated as (out-degree − in-degree)/(out-degree + in-degree). crTFs with hierarchy scores from 0.33 to 1 were classified as top roles, those with scores between −0.33 and 0.33 were classified as middle roles, and those with scores from −1 to −0.33 were classified as bottom roles. The table lists the in-degree, out-degree, hierarchy score, and hierarchical role of each crTF in PV, MZ, and CV hepatocytes, and compares the conservation of hierarchical roles among shared crTFs across the three states.

**Supplemental Table S9. Conserved shared crTFs across PV, MZ, and CV hepatocytes.**

Worksheet 1 lists shared crTFs that maintained both invariant core–periphery assignments and conserved hierarchical levels across the three states. Worksheet 2 lists the number of invariant core and invariant peripheral crTFs at each hierarchical position.

**Supplemental Table S10. Liver-disease associations of broadly expressed upstream-core crTFs.**

Expression breadth from GTEx bulk RNA-seq across 42 human tissues. Disease associations and dedifferentiation/identity links from the cited literature.

**Supplemental Table S11. Classification of published cell-type-level hepatocyte CRN factors in the state-level CRNs.**

Worksheet 1 lists factors of the Dubois-Chevalier core circuitry (CoRC, n = 13) and extended network (CONNECT, n = 26), classified as invariant (members of the 143-crTF invariant sub-network), shared non-invariant (present in all three states but shifting position), two-state, or absent. Worksheet 2 lists the presence of the 12 upstream-core crTFs in the core regulatory networks of Saint-André et al. (their Supplemental Table S3, 84 cell and tissue types); 8 of 12 appear in at least one network.
